# Robust and High-Throughput Analytical Flow Proteomics Analysis of Cynomolgus Monkey and Human Matrices with Zeno SWATH Data Independent Acquisition

**DOI:** 10.1101/2022.11.30.518440

**Authors:** Weiwen Sun, Yuan Lin, Yue Huang, Josolyn Chan, Sonia Terrillon, Anton I. Rosenbaum, Kévin Contrepois

## Abstract

Modern mass spectrometers routinely allow deep proteome coverage in a single experiment. These methods are typically operated at nano and micro flow regimes, but they often lack throughput and chromatographic robustness, which is critical for large-scale studies. In this context, we have developed, optimized and benchmarked LC-MS methods combining the robustness and throughput of analytical flow chromatography with the added sensitivity provided by the Zeno trap across a wide range of cynomolgus monkey and human matrices of interest for toxicological studies and clinical biomarker discovery. SWATH data independent acquisition (DIA) experiments with Zeno trap activated (Zeno SWATH DIA) provided a clear advantage over conventional SWATH DIA in all sample types tested with improved sensitivity, quantitative robustness and signal linearity as well as increased protein coverage by up to 9-fold. Using a 10-min gradient chromatography, up to 3,300 proteins were identified in tissues at 2 µg peptide load. Importantly, the performance gains with Zeno SWATH translated into better biological pathway representation and improved the ability to identify dysregulated proteins and pathways associated with two metabolic diseases in human plasma. Finally, we demonstrate that this method is highly stable over time with the acquisition of reliable data over the injection of 1,000+ samples (14.2 days of uninterrupted acquisition) without the need for human intervention or normalization. Altogether, Zeno SWATH DIA methodology allows fast, sensitive and robust proteomic workflows using analytical flow and is amenable to large-scale studies. This work provides detailed method performance assessment on a variety of relevant biological matrices and serves as a valuable resource for the proteomics community.

## INTRODUCTION

Recent advances in mass spectrometry (MS) based proteomics including instrumentation, chromatography, data acquisition strategies and processing software allow high quality deep proteome coverage in a single-run experiment. These advancements are fueling a growing interest in utilizing those technologies in applications requiring high throughput phenotypic readouts at scale. Notably, MS-based proteomics is emerging as a key player in drug discovery with the promise to accelerate i) target identification and validation, ii) lead discovery and optimization as well as iii) pre-clinical and clinical assessments of drug safety and efficacy [1]. Recently, deep proteome profiling of ∼900 cancer cell lines by MS identified thousands of potential biomarkers of cancer vulnerabilities [2]. In addition, proteomics has been widely applied to the field of precision medicine enabling patient stratification [3] and improving patient management [4].

Among discovery-driven acquisition strategies, data-dependent acquisition (DDA) and data-independent acquisition (DIA) methodologies are the most popular. While in DDA a predefined number of the most abundant peptides from survey scans are selected for fragmentation, all peptides in a sliding m/z window in MS1 scans are selected for fragmentation in DIA. Therefore, DIA demonstrated better quantification robustness, data completeness and sensitivity than DDA [5]. Among DIA methodologies, Sequential Windowed Acquisition of All Theoretical Fragment Ion Mass Spectra (SWATH-MS) is a widely used methodology that allows deep proteome coverage with quantitative consistency and accuracy [6]. A recent multi-laboratory evaluation study demonstrated its reproducibility as well as quantitative and qualitative performances [7].

To date, SWATH-MS methods have been primarily operated at nano [7] and micro flow regime [2] providing excellent proteome depth and data quality. However, these methods often lack throughput and chromatographic robustness necessary for the analysis of large studies. In this context, we developed, optimized and benchmarked robust and high-throughput analytical flow LC-MS methods with Zeno SWATH DIA approach. Taking advantage of the Zeno trap, Zeno SWATH methods consistently provided higher sensitivity and protein coverage as well as superior data quality across a wide variety of biological matrices in comparison to SWATH. Furthermore, Zeno SWATH DIA improved pathway analysis and enhanced the ability to identify plasma biomarkers for two metabolic diseases in humans, demonstrating the clinical application of this novel method.

## RESULTS

### Zeno Trap Enhances Sensitivity and Proteome Coverage

The Zeno trap featured on SCIEX 7600 ZenoTOF instrument is a linear ion trap located after the collision cell that reportedly improves detectability of fragments by up to 20-fold [8-10]. This increased sensitivity can mitigate the main drawback of analytical flow methods (*i*.*e*. limited sensitivity) in comparison to more conventional micro or nano flow set-ups. To assess the gain in information obtained with Zeno trap activated, we prepared protein extracts from a variety of biological matrices including ten cynomolgus monkey tissues (stomach, kidney, spleen, lung, colon, liver, rectum, cecum, bone marrow and heart), three human biofluids (sputum, bronchoalveolar lavage [BAL] fluid and non-depleted plasma) and two *in vitro* samples from air-liquid interface cultures (mucus and media) (**Fig. 1A**).

**Figure 1.**
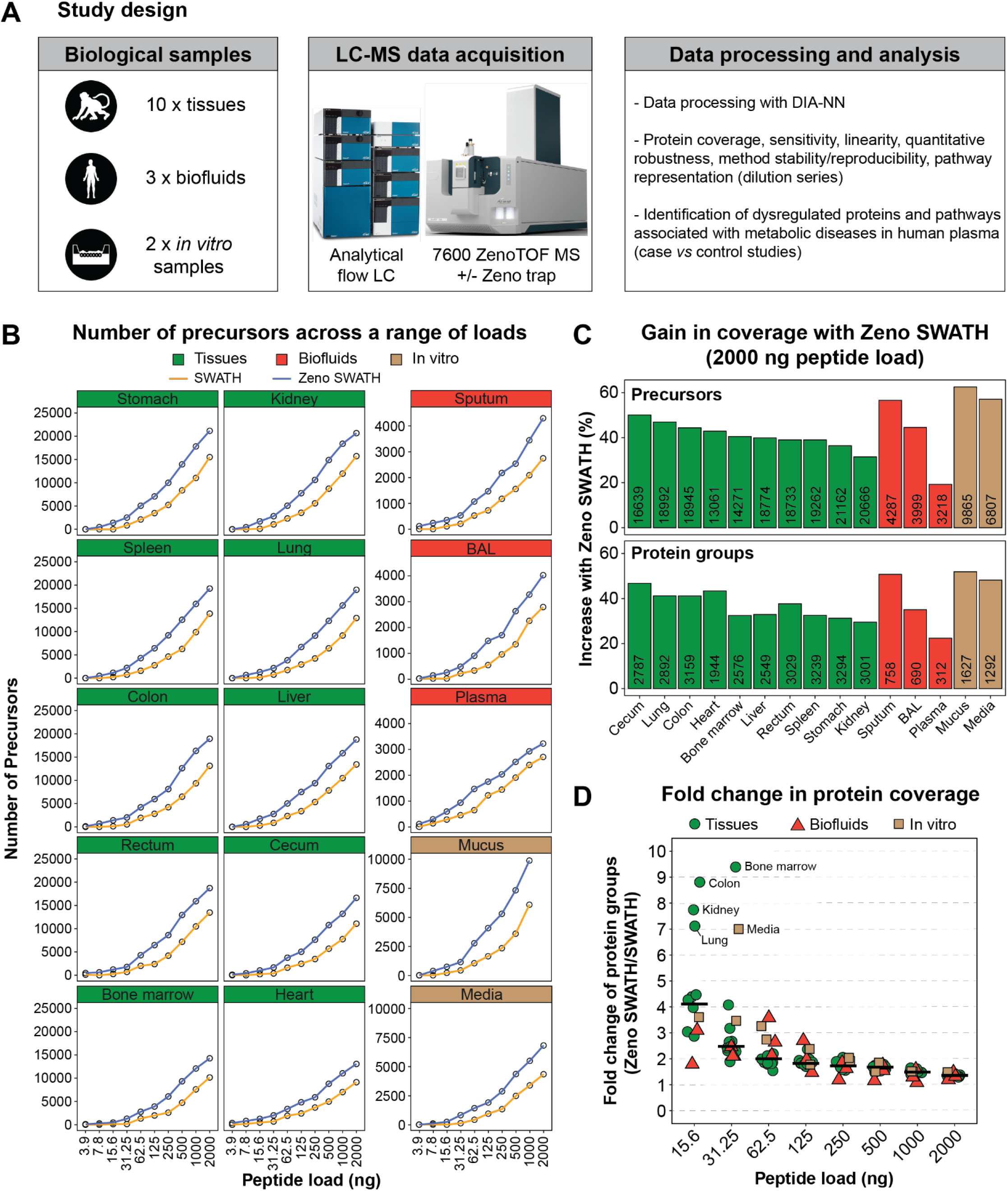
Study design, assay sensitivity and proteome coverage. (**A**) Overview of the study design including cynomolgus monkey and human matrices analyzed with an analytical flow SWATH DIA workflow with and without Zeno trap activated and analysis plan. (**B**) Number of precursors identified across technical triplicates in various biological matrices at different peptide loads in SWATH and Zeno SWATH DIA modes. Peptide loads range from 3.9 ng to 2000 ng for all sample types except for mucus that ranges from 3.9 ng to 1000 ng. (**C**) Gain in precursors and protein groups with Zeno SWATH in comparison to SWATH at the highest peptide load (2000 ng). The number of precursors and protein groups detected with Zeno trap activated is indicated in bars. (**D**) Fold change in protein groups identified in Zeno SWATH mode across peptide loads. The horizontal bars represent the median. See also **Figure S1**.

Peptide separation was performed at an analytical flow regime with a 10-min gradient chromatography to ensure throughput and robustness and LC-MS parameters were optimized to identify the most protein groups and precursors in each biological matrix. The parameters evaluated included flow rate, source parameters, MS1 and MS2 accumulation time as well as SWATH window number and window width (**Methods** and **Tables S1** & **S2**). Notably, the optimal parameters selected were slightly different for each sample type reflecting variable protein compositions and expression levels.

Sensitivity and protein coverage were determined on ten-point dilution series across all biological matrices (3.9 ng to 2000 ng) using optimized LC-MS methods with and without Zeno trap activated and raw data were processed using *in silico* predicted libraries in DIA-NN (**Methods**) [11]. Zeno SWATH DIA methods systematically detected more precursors and protein groups at all peptide loads across all sample types in comparison to conventional SWATH DIA methods run on the same instrument (**Fig. 1B** & **S1A**). At maximum load (2000 ng), enabling Zeno trap increased the number of detected precursors by 20-60% and protein groups by 20-50% (**Fig. 1C**). Among the various sample types, non-depleted plasma benefited the least from the boost in sensitivity while sputum, mucus and media benefited the most. At the 2000 ng load, Zeno SWATH methods detected 13100 to 21200, 3200 to 4300 and 6800 to 9900 precursors in tissues, biofluids and *in vitro* samples, respectively corresponding to 1950 to 3300, 300 to 750 and 1300 to 1600 protein groups.

While Zeno trap improved protein coverage at high loads, the gain in sensitivity was the most remarkable at low peptide loads (15.6 ng and 31.25 ng) with increases in protein and precursor coverage by 2-4 fold across the various matrices and up to 9-fold for bone marrow and colon (**Fig. 1D** & **Fig. S1B**). Tissues and in particular bone marrow, colon, kidney and lung benefited the most from the increase in sensitivity. Finally, while SWATH methods were unable to detect any precursor at very low peptide loads (3.9 ng and 7.8 ng), Zeno SWATH methods typically enabled detection of precursors and protein groups at these loads demonstrating the utility of this technology when sample input is limited (**Fig. 1B** & **Fig. S1A**).

### Zeno Trap Improves Pathway Coverage

First, we investigated global protein expression profiles of cynomolgus monkey tissues using hierarchical clustering to assess data quality. As expected, samples from the gastrointestinal tract (cecum, rectum, colon and stomach) and hematopoietic tissues (bone marrow and spleen) clustered together indicating similarities in protein expression (**Fig. S2**). These results confirm previous observations made in human tissues [12].

Next, we asked whether the gain in protein coverage with Zeno trap activated could improve pathway representation. Pathway enrichment analysis was performed using Gene Ontology (GO) database [13, 14] and revealed a systematic increase in pathway coverage across GO categories and peptide loads with larger gains at lower loads (**Fig. 2** & **Table S3**). We observed an average increase in GO Process, Function and Component pathways by 159%, 98% and 133%, respectively at 31.25 ng peptide load in tissues. Similarly, pathway coverage increased by 179% and 181% on average across GO databases in biofluids and *in vitro* samples, respectively at 31.25 ng peptide load. While the gain in pathway coverage was mostly comparable among tissues and *in vitro* samples, results from biofluid specimens were more variable. In particular, pathway coverage increase in plasma was modest in comparison to other biofluids (54% on average across GO databases at 31.25 ng peptide load) reflecting the limited improvement in the number of proteins detected. Altogether, these results demonstrate that Zeno SWATH DIA provides more biologically meaningful information by covering more pathways.

**Figure 2.**
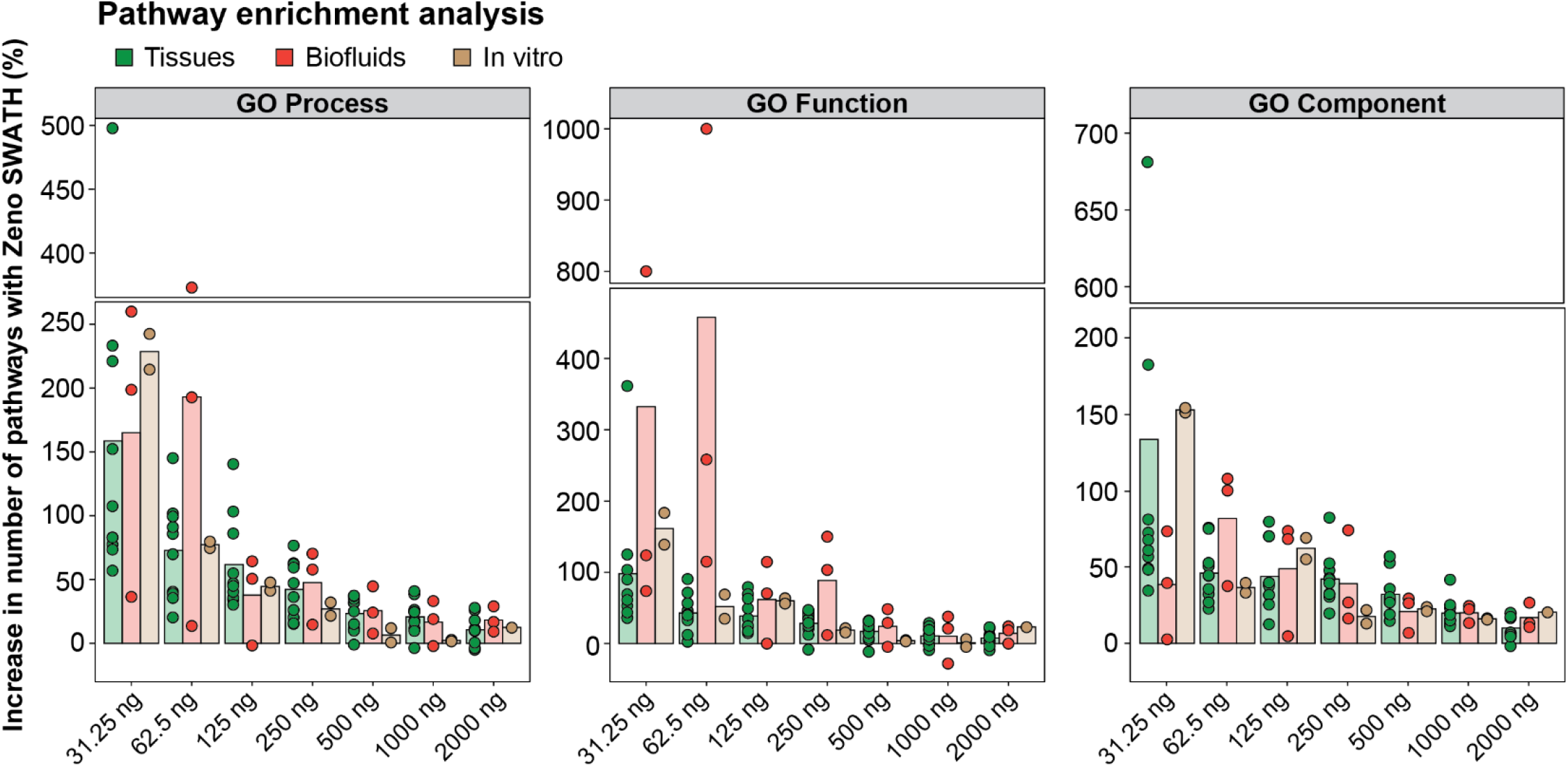
Pathway representation with and without Zeno trap activated. Gain in pathway coverage using proteins identified in Zeno SWATH DIA mode in comparison to SWATH DIA. Bars represent the mean across samples belonging to each matrix type (tissues [n=10], biofluids [n=3], *in vitro* samples [n=2]). Pathway enrichment was performed using Gene Ontology database. See also **Figure S2**.

### Zeno Trap Enhances Quantitative Robustness and Signal Linearity and Method Performance is Stable Over Time

In addition to proteome coverage and sensitivity, the quantitative quality of the data generated in Zeno SWATH mode was assessed leveraging dilution series and technical triplicates at each level. Quantitative robustness -as calculated by the proportion of proteins robustly quantified with a CV below 0.2 across triplicate injections – was significantly improved in tissues at all peptide loads except at 31.25 ng (**Fig. 3A**). For instance, 91.3% ± 1.3% and 86.5% ± 2.1% of protein groups were detected with a CV < 0.2 in Zeno SWATH and SWATH modes, respectively at 2000 ng peptide load. Similar observations were made in biofluids and *in vitro* samples (**Fig. S3A**). Signal linearity was assessed by calculating Pearson coefficients of correlation of precursor intensities against peptide loads. While high coefficients of correlation were observed in both modes (r > 0.98 in average), Zeno SWATH significantly improved linearity in all matrices (P-value < 1.3E-04, **Fig. 3B** & **Fig. S3B**).

**Figure 3.**
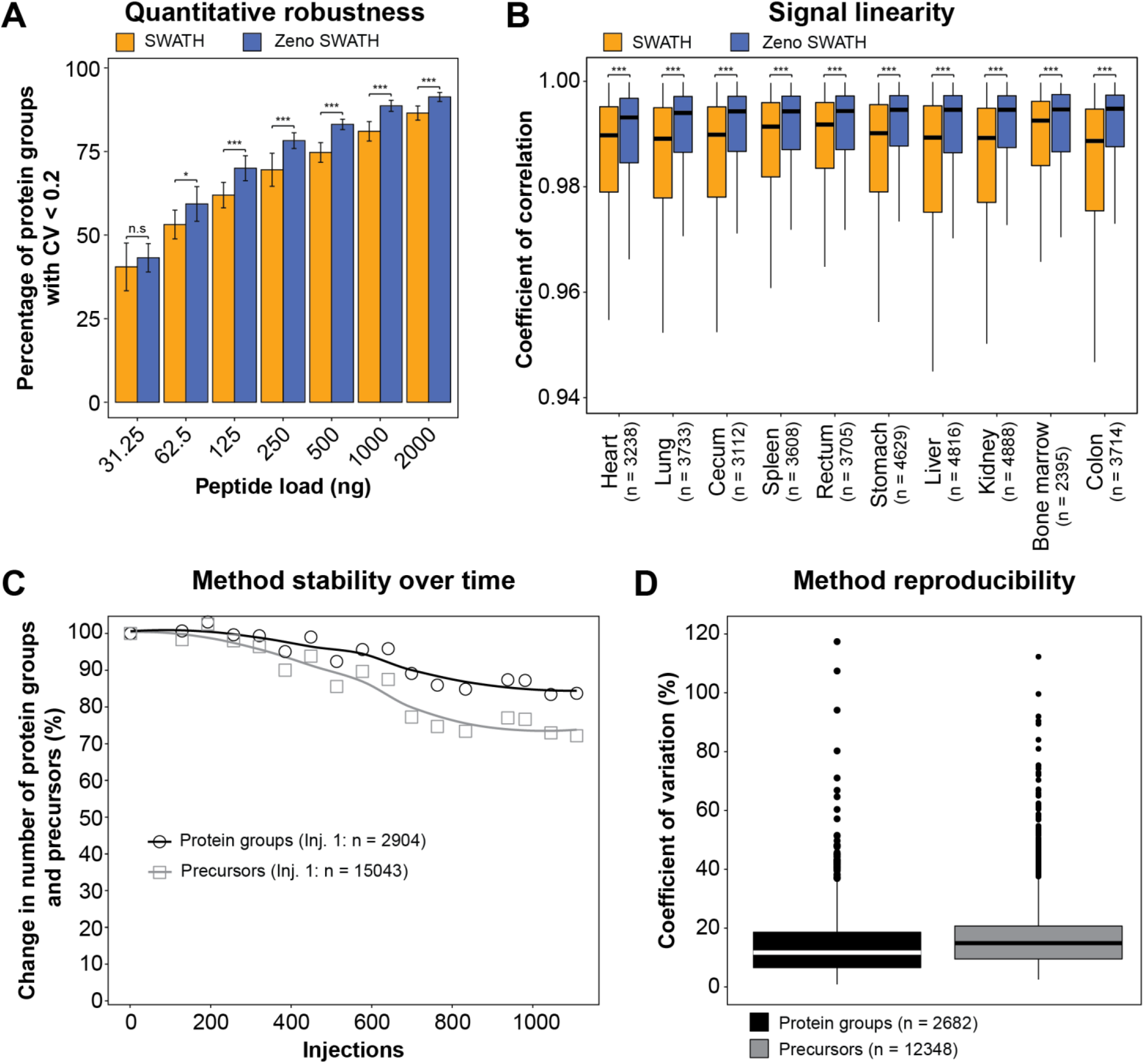
Evaluations of quantitative robustness, signal linearity and method stability over time. (**A**) Proportion of proteins with a coefficient of variation (CV) below 0.2 calculated across technical triplicates at different loads in SWATH and Zeno SWATH DIA modes. Only proteins detected in 100% of the triplicates were considered and only conditions with more than 50 proteins identified were plotted. Bars represent the mean across 10 cynomolgus monkey tissues and the error bars represent the standard deviation. A two-sided Welch’s t-test was used for differential analysis. n.s: not significant, *: P-value < 0.05, **: P-value < 0.01, *** P-value < 0.001. (**B**) Boxplot showing the distribution of Pearson coefficients of correlation of precursor abundances against peptide loads for cynomolgus monkey tissues. Only precursors detected in at least 3 levels in both SWATH and Zeno SWATH DIA modes were considered in the analysis. The number of precursors is indicated in parenthesis. The box illustrates the first and third quartile, the whiskers are 1.5 times the interquartile range and the horizontal bar depicts the median of the distribution. A two-sided Mann–Whitney U test was used for differential analysis. *** P-value < 0.001. (**C**) Number of protein groups and precursors detected in quality control HeLa protein digest samples (n = 17) in Zeno SWATH DIA mode along the study. The curves represent a LOESS (locally estimated scatterplot smoothing) fit. The numbers of protein groups and precursors detected in the first injection of HeLa digest are indicated in parenthesis. (**D**) Coefficient of variation of protein groups and precursors detected in at least half of the HeLa samples. The number of protein groups and precursors considered for the analysis are indicated in parenthesis. The box illustrates the first and third quartile, the whiskers are 1.5 times the interquartile range and the horizontal bar depicts the median of the distribution. See also **Figure S3**.

We demonstrated that the combination of analytical flow chromatography with a 10-min gradient and Zeno SWATH DIA enhanced sensitivity, expanded protein coverage, and generated higher quality data. However, for this method to be well suited to large-scale studies, method stability over time remained to be shown. To evaluate method stability, HeLa digests (n=17) were injected repetitively along the study which consisted in uninterrupted injections of 1,000+ samples (equivalent to ∼500 µg peptides injected in total) that lasted 14.2 days. We observed a modest decrease in the number of precursors (27.8%) and protein groups detected (16.6%) demonstrating method robustness (**Fig. 3C**). In addition, signal intensity remained stable without the need for normalization with median coefficients of variation across all HeLa digest injections of 14.9% and 11.7% for precursors and protein groups, respectively (**Fig. 3D**).

Altogether, we have demonstrated robust and consistent performance of our high throughput LC-MS method over the injection of 1,000+ samples without the need for human intervention (*e*.*g*., instrument maintenance, tuning or cleaning) and data normalization.

### Zeno Trap Identifies More Potential Disease Biomarkers in Plasma Samples

Finally, we assessed the ability of Zeno SWATH to identify biomarkers and inform our understanding of disease mechanisms. In this context, we profiled protein content in plasma from patients diagnosed with non-alcoholic steatohepatitis (NASH) and type 2 diabetes (T2D) with and without Zeno trap activated. Subject demographics information can be found in **Table S4**.

Differential analysis of samples from NASH patients *vs* healthy individuals identified 24 and 44 significantly dysregulated proteins in SWATH and Zeno SWATH modes, respectively (FDR < 0.1, **Fig. 4A, Table S5**). As expected, most significant proteins in SWATH mode (87.5%) were also significant with Zeno SWATH and included proteins that have been associated with NASH onset (leucine-rich alpha-2-glycoprotein 1 [LRG1] [15]), progression (inter-alpha-trypsin inhibitor heavy chain 4 [ITIH4] [16]) and severity (alpha-1 antitrypsin [SERPINA1] [17]) as well as fibrosis (extracellular matrix protein 1 [ECM1] [18]). Zeno SWATH identified additional dysregulated proteins among which 36.4% were not significant and 15.9% were not detected in SWATH mode (**Fig. 4C**). Some of these proteins were previously described in the context of NASH and included oxidative stress marker glutathione peroxidase 3 (GPX3) [19] and coagulation factors F9 and F10 [20]. Pathway enrichment analysis identified expected pathways associated with metabolism and liver inflammation such as ‘LXR and FXR activation’ [21] and ‘acute phase response signaling’ [22] as well as coagulation [20] (*i*.*e*. ‘coagulation system’) (**Table S6**). While these pathways were significant in both SWATH and Zeno SWATH modes, their significance was enhanced with Zeno trap activated (**Fig. 4D**). In particular, the significance of ‘coagulation system’ improved dramatically (P-value = 3.1E-02 in SWATH mode and P-value = 7.9E-16 in Zeno SWATH mode) indicating that additional significant proteins in Zeno SWATH mode belong to this pathway.

**Figure 4.**
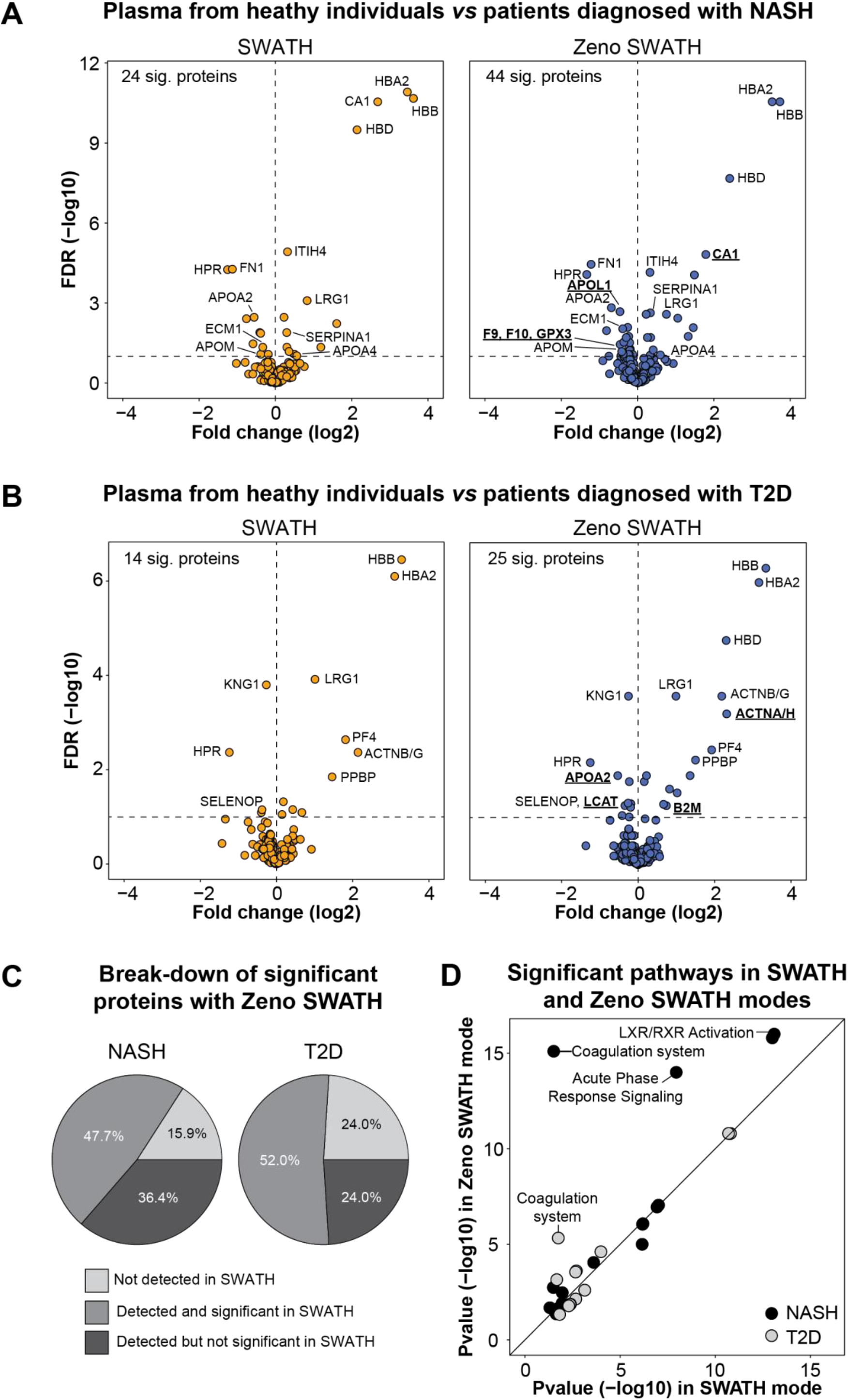
Clinical proteomics biomarker discovery. Volcano plots representing differential proteins in individuals diagnosed with non-alcoholic steatohepatitis (NASH, n = 20) (**A**) and type 2 diabetes (T2D, n = 10) (**B**) in comparison to healthy individuals (n = 20). A two-sided Welch’s t-test was used for differential analysis and proteins with FDR < 0.1 were considered significant. Proteins underlined and in bold are uniquely significant in Zeno SWATH mode. (**C**) Pie charts representing the proportion of significant proteins (FDR < 0.1) in Zeno SWATH DIA mode that were i) not detected, ii) detected and significant, and iii) detected but not significant in SWATH DIA mode. (**D**) Comparison of pathway significance using significant proteins identified in SWATH and Zeno SWATH DIA modes in NASH and T2D studies. Ingenuity Pathway Analysis was used for enrichment analysis and pathways with P-values < 0.05 were considered significant.

Similarly, 14 and 25 differential proteins were identified in plasma from patients with T2D *vs* healthy individuals in SWATH and Zeno SWATH modes, respectively (FDR < 0.1, **Fig. 4B, Table S5**). Among overlapping proteins, pro-platelet basic protein (PPBP) has been associated with diabetic nephropathy podocyte injury [23], selenoprotein P (SELENOP) is a hepatokine that causes insulin resistance in T2D [24] and platelet factor 4 (PF4/CXCL4) reflects platelet function [25]. Zeno SWATH identified additional relevant proteins involved in the pathophysiology of T2D such as beta-2 microglobulin (B2M) - a potential biomarker of diabetic nephropathy [26] and lecithin–cholesterol acyltransferase (LCAT) - a marker of metabolic syndrome and diabetes mellitus [27]. Among significant proteins with Zeno SWATH, 24.0% were not significant and 24.0% were not detected in SWATH mode (**Fig. 4C**). At the pathway level, results were similar with and without Zeno trap activated except pathway significance improved for a subset of them including ‘coagulation system’ (P-value = 1.8E-02 in SWATH mode and P-value = 4.7E-6 in Zeno SWATH mode) (**Fig. 4D, Table S6**) [28].

Altogether, Zeno trap resulted in identifying ∼80% more differential proteins in plasma from patients diagnosed with metabolic diseases and these proteins and associated pathways were relevant to each disease condition.

## DISCUSSION

In this work, we developed and optimized analytical flow LC-MS methods with Zeno SWATH DIA across a range of biological matrices and assessed the performance gain relative to SWATH DIA methods. Enabling the Zeno trap systematically improved all the performance metrics examined encompassing coverage in proteins and pathways, data quality and ability to identify plasma biomarkers in clinical samples.

MS-based proteomics is rapidly evolving owing to incremental hardware and software improvements [6, 11, 29-32]. As instruments and acquisition methods are becoming more sensitive and processing software more elaborated, the proteomics community is moving towards shorter gradients and higher flow rates to enable wider application of the methodology to studies requiring large sample size. One example uses a 5-min gradient method at 800 µL/min to profile plasma proteins [33]. While such a method is beneficial for plasma analysis due to its unique distribution in protein abundance, protein coverage in other sample types remains limited due to decreased sensitivity. The gain in sensitivity provided by the Zeno trap improved protein and precursor coverage by up to 50% at 2 µg peptide load with a 10-min gradient analytical flow chromatography. These methods detected up to 3,300 proteins from tissues which is close to performances obtained at micro flow using slightly longer gradients [34]. The gain in protein coverage was the most notable at low peptide loads with up to 9-fold increase highlighting the utility of the technology when sample input is limited.

With omics technologies becoming more mature, they are being increasingly used in epidemiological studies generating high content quality datasets for disease risk prediction and prevention as well as disease mechanism investigation. Focusing on circulating blood markers, metabolomics and proteomics are the two most popular approaches and successfully generated predictive models for a wide range of disorders including metabolic, cardiovascular and neurological diseases [35-38]. Even though proteomics has been successful in capturing important aspects of human disease pathophysiology, adding information from complementary omics layers has improved sensitivity and specificity of predictive models in studies of complex diseases (*i*.*e*. type 2 diabetes) [39], and physiological processes (*i*.*e*. acute exercise and pregnancy) [40, 41]. In addition, proteomics has the potential to benefit interventional studies with the discovery of safety and efficacy biomarkers as well as the investigation of mechanisms of drug toxicity and drug action. For instance, large scale plasma proteomics revealed mechanisms by which torcetrapib - a cholesterol ester transfer protein inhibitor that was being developed to treat hypercholesterolemia and prevent cardiovascular disease – may exert harmful effects [42]. The authors were also able to identify a 9-protein signature that could predict an increase in derived risk from torcetrapib within just 3 months of treatment.

In order to further democratize the use of proteomics technologies in clinical trials, there is a clear need to develop high throughput and robust methods that can capture the proteome in-depth while minimizing the need for instrument maintenance. Analytical flow regime provides chromatographic robustness and signal stability over time as demonstrated by the modest decrease in sensitivity after the uninterrupted injection of 1000+ samples and high reproducibility of quality control samples injected throughout the study. Even though post-acquisition methods have been developed to improve instrument reproducibility over time, they are typically computationally intensive and imperfect in deconvoluting unwanted variation [43, 44].

In conclusion, our comparative study reveals that SWATH DIA with Zeno trap enabled improves sensitivity, quantitative robustness and signal linearity and expands protein coverage in all sample types tested. The gain in protein coverage translated into better biological pathway coverage and enabled identification of potential disease biomarkers and dysregulated pathways in a cross sectional setting. In addition, the methods developed using analytical flow chromatography were extremely robust minimizing the need for human intervention in large-scale studies. Thus, Zeno SWATH DIA technology is well suited for applications that require a phenotypic readout at scale and will likely benefit drug development, toxicological, mechanistic and clinical biomarker discovery studies.

## EXPERIMENTAL PROCEDURES

### Experimental Design and Statistical Rationale

The following experiments were performed: i) triplicate injections of ten-point dilution series from ten cynomolgus monkey tissues, three human biofluids and two *in vitro* samples from air-liquid interface cultures, ii) singlicate injection of clinical plasma samples from twenty healthy individuals and twenty individuals diagnosed with non-alcoholic steatohepatitis (NASH), and ten individuals diagnosed with type 2 diabetes (T2D), and iii) repetitive singlicate injection of commercial HeLa cell protein digest along the sequence (n = 17). All experiments were conducted in SWATH DIA mode with and without Zeno trap activated on the same instrument and HeLa experiments were performed with Zeno trap activated. All data were acquired in a single uninterrupted sequence. Dilution series were acquired to assess the performance of SWATH DIA experiments with Zeno trap activated (*i*.*e*. sensitivity, coverage, signal linearity) and technical replicates were used to evaluate quantitative robustness by calculating coefficients of variation (CVs). Dilution series were run from low to high loads to avoid potential carryover. Clinical plasma samples were run in a random order. Repetitive injections of HeLa extracts were used to evaluate instrument stability over time and method reproducibility. Data processing was performed with DIA-NN (v1.8.1). Additional information on the samples analyzed, data acquisition and processing procedures can be found below.

### Materials and Reagents

Acetonitrile (ACN, LC-MS grade), water (LC-MS grade), methanol (LC-MS grade), 2-chloroacetamide, phosphoric acid solution, and sodium dodecyl sulfate (SDS) were purchased from Sigma-Aldrich. Bond-Breaker tris(2-carboxyethyl)phosphine (TCEP) solution, 1M triethylammonium bicarbonate (TEAB), trypsin/Lys-C protease mix (cat# A40009), Pierce protease inhibitor mini tablets (cat# A32953), Pierce HeLa protein digest standard (cat# 88329), and formic acid (LC-MS grade) were obtained from Thermo Fisher Scientific.

### Sample Collection and Origin

Cynomolgus monkey tissue samples including bone marrow, cecum, colon, heart, kidney, liver (left lateral), lung, rectum, spleen, and stomach were collected from healthy cynomolgus monkeys immediately after sacrifice. The procedures involving the care and use of animals was reviewed and approved by Charles River - Nevada Institutional Animal Care and Use Committee (IACUC) and the care and use of animals were conducted with guidance from the guidelines of the USA National Research Council. Experiments were conducted using a single tissue piece from a single specimen. Bronchoalveolar lavage (BAL) fluids were collected from 10 healthy individuals using sterile saline. Samples were then filtered with 100 µm filters and centrifuged at 1500 rpm for 10 minutes at 4°C. The resulting supernatants were immediately aliquoted and frozen at −80°C until further analysis. A pool of all the BAL samples was used in this work. BAL collection was approved by The National Jewish Health institute and informed consent forms were obtained from all participants. Commercial human EDTA-plasma samples from 20 individuals self-reported as healthy were purchased from BioIVT and EDTA-plasma samples from 20 individuals diagnosed with NASH, and 10 individuals diagnosed with T2D were purchased from ProteoGenex (lot numbers and subject demographics information are available in **Table S4**). A pool of all healthy plasma samples was used for the dilution series experiment. Induced human sputum samples from two healthy donors were purchased from BioIVT and pooled for analysis. Mucus and media samples were obtained from air-liquid interface (ALI) *in vitro* bronchial cell cultures and pooled from experiments performed with primary bronchial cells from 3 to 6 human donors. Cryopreserved primary normal human bronchial epithelial (NHBE) cells were purchased from Lonza (CC-2540 & CC-2540S) and ATCC (PCS-300-010). NHBE cells were expanded and maintained at an ALI as described previously [45, 46] with minor modifications. Briefly, NHBE cells were expanded in BEGM growth medium (Lonza #CC-3170) before being plated onto Collagen type IV (cat# C7521, Sigma)-coated 0.4-μm pore-size transwells (cat# CLS3470, Corning) and cultured under liquid-liquid interface growth conditions with complete BEGM medium. ALI culture conditions were initiated when cells reached confluence by removal of apical medium and replacement of basolateral growth medium with B-ALI differentiation medium (cat# 193517, Lonza) supplemented with SingleQuots (cat# 193515, Lonza) and retinoic acid (cat# #R2625, Sigma). After 4 weeks of airlift culture, the mucus overlying the epithelium was washed apically using PBS before being collected along with medium samples. All the samples were stored at −80°C prior to the study.

### Sample Preparation

Approximately 25 mg of frozen tissues were transferred to 2 mL Eppendorf Protein Lobind tubes containing one 5 mm stainless steel bead (cat# 69989, Qiagen) and 500 µL of lysis buffer consisting of 5% SDS in 50 mM TEAB with protease inhibitors cocktail (cat# A32953, Thermo Fisher Scientific). Tissue pieces were disrupted with TissueLyser LT (Qiagen) at 50 Hz (1 min ‘on’, 1 min ‘off’ on ice) until fully homogenized. Four microliters of plasma was added to 46 µL of lysis buffer and sputum (200 µL), BAL (300 µL), mucus (300 µL) and media (150 µL) samples were first dried to completion under a stream of nitrogen (TurboVap 96, Biotage) and resolubilized in 50 µL of lysis buffer. Homogenates were sonicated on ice for 5 min at 30% amplitude (1 min ‘on’, 15s ‘off’) with a Qsonica sonicator (Q700MPX) and centrifuged for 10 min at 14,000g at 18°C. The supernatants were then collected, and protein concentrations were determined using Pierce BCA Protein Assay Kit (cat# 23225, Thermo Fisher Scientific) following the manufacturer’s instructions. Reduction and alkylation of proteins were performed after incubation of the extracts at 95°C for 10 min by adding 5 µL of 100 mM TCEP and 5 µL of 400 mM chloroacetamide each followed by an incubation at 95°C for 5 min.

On-column digestion was performed using S-Trap sample processing technology in 96-well plate format (100 – 300 μg per well, cat# NC1508276, Protifi LLC) following the manufacturer’s instructions. First, proteins were further denatured by adding 5 µL of phosphoric acid (12%) followed by 300 µL of S-Trap binding buffer (90% MeOH, 100 mM final TEAB, pH 7.1) to protein extracts (15-150 µg of proteins). Proteins were then loaded and captured on the S-Trap columns following centrifugation at 1,500g for 2 min at room temperature. Proteins were then washed 3 times with 200 μL S-Trap binding buffer (90% MeOH, 100 mM final TEAB, pH 7.1) and centrifuged at 1,500g for 2 min. Protein digestion was performed overnight at 37°C with trypsin/Lys-C protease mix (cat# A40009, Thermo Fisher Scientific) in 125 µL 50 mM TEAB (1:30 enzyme:protein ratio). Stepwise elution of peptides was performed with 80 µL of 50 mM TEAB, 80 µL of 0.2% formic acid in water, and 80 µL of 0.2% formic acid in 50% ACN each followed by centrifugation at 1,500g for 2 min. Flowthroughs were then pooled and dried under a stream of nitrogen (TurboVap 96, Biotage) and peptides were resuspended in 100 µL of 0.1% (v/v) formic acid in 5% (v/v) ACN. Peptide concentrations were determined using Pierce Quantitative Colorimetric Peptide Assay (cat# 23275, Thermo Fisher Scientific), and samples were frozen at −80°C until LC-MS data acquisition.

### Untargeted Proteomics by LC-MS

Peptide extracts were analyzed using an Exion LC system coupled with a 7600 ZenoTOF mass spectrometer (SCIEX). Peptides were separated on an Acquity BEH C18 column 2.1 × 150 mm, 1.7 μm (cat# 186003687, Waters) and mobile phase solvents consisted in 0.1% formic acid in water (A) and 0.1% formic acid in ACN (B). Peptides were eluted from the column using a 10-minute linear gradient (5-40% B) at a flow rate of 0.2 mL/min and the oven temperature was set at 40°C. Autosampler temperature was set at 10°C. The 7600 ZenoTOF was equipped with an OptiFlow Turbo V ion source and operated in SWATH mode with or without Zeno trap activated. The source conditions were as follows: ionization voltage: 5500 V, positive polarity, temperature: 500°C, ion source gas 1: 60 psi, ion source gas 2: 80 psi, curtain gas: 40 psi. The acquisition settings were as follows: number of Q1 windows: 50, MS1 accumulation time: 100 ms, MS1 *m/z* range: 400–900 Da, MS2 accumulation time: 18 ms (plasma and media) or 15 ms (rest of the samples), MS2 *m/z* range: 100–900 Da. Q1 window widths were determined for each sample type from DDA runs using SWATH Variable Window Calculator_V1.2 provided by SCIEX. All tissues were run with the same method optimized on liver sample. External calibration was performed every 3 to 8 runs using the automatic calibration function and ESI positive calibration solution for SCIEX X500B system (cat# 5049910, SCIEX). Ten-point dilution series (3.9 ng – 2000 ng) were prepared the day prior to the experiment, stored at 4°C overnight and injected from low to high concentrations in triplicates in both SWATH DIA and Zeno SWATH DIA modes. Healthy and diseased plasma samples were injected in singlicate in a random order in SWATH mode first followed by Zeno SWATH. All samples were injected no more than 1.7 days after being loaded in the autosampler. Signal stability over time and method robustness were monitored by repetitive injection of 800 ng of HeLa protein digest (cat# 88329, Thermo Fisher Scientific) in Zeno SWATH mode. HeLa protein digest sample was left in the autosampler and the detector parameters were left unchanged during the whole duration of the study.

### Data Processing

All raw data were processed with DIA-NN (v1.8.1) with a fragment ion *m/z* range of 200–1800, automatic settings of mass accuracy at the MS2 and MS1 level and scan window, protein inference of ‘Genes’ and quantification strategy of “robust LC (high precision)”. For dilution series, triplicate samples from each dilution level and acquisition mode were processed separately with cross-run normalization disabled and analyzed with match-between-runs (MBR) enabled. Clinical plasma samples from each case study and each acquisition mode were processed separately with cross-run normalization enabled (‘RT-dependent’) and analyzed with match-between-runs (MBR) enabled. All database searches were performed using *in silico* predicted libraries from Human UniProt canonical sequence database (UP000005640) (human biofluids and *in vitro* samples) and UniProt canonical sequence database (UP000233100) (cynomolgus monkey tissues). Predicted libraries were generated using the following parameters: Trypsin/P, one missed cleavage allowed, N-term M excision, C carbamidomethylation, peptide length range: 7-30 amino acids, precursor charge range: 1-4. Details on library of decoy precursors generation and data matching to spectral libraries can be found in the original DIA-NN article [11].

### Data Analysis and Visualization

Unless otherwise mentioned, analyses were performed in R (v4.1.3) and RStudio (v1.4.1103). For all analyses pertaining to dilution series, global q-value thresholds of 1% and 5% were used for precursors and protein groups, respectively (**Fig. 1, 2, 3, S1, S2** & **S3**). Analysis of clinical plasma samples was performed using global q-values thresholds of 1% for precursors and protein groups (**Fig. 4**).

Protein content similarity in tissue samples was assessed by calculating pairwise Pearson correlations of median expression levels across triplicates using protein groups present in at least 50% of the tissue samples (**Fig. S2**). The heatmap was plotted using Pheatmap package (v1.0.12) in R and the clustering distance used was ‘maximum’. Pathway analysis in tissues was performed using STRINGdb package (v2.6.5) in R and database version 11.5 (**Fig. 2**). Pathways with FDR < 0.05 were considered significant. Coefficients of variation (CVs) were calculated across triplicate injections and the proportions of protein with a CV < 0.2 were determined relative to proteins detected in 100% of the triplicates (**Fig. 3A** & **S3A**). Differential analysis was performed using a two-sided Welch’s t-test. Pearson coefficients of correlation of precursor abundances against peptide loads were calculated in all matrices (**Fig. 3B** & **S3B**). Only precursors detected in at least 3 levels in both SWATH and Zeno SWATH modes were considered in the analysis. A two-sided Mann–Whitney U test was used for differential analysis. Coefficients of variation in HeLa protein digests were calculated using protein groups and precursors detected in at least half of the samples (**Fig. 3D**). Analysis of plasma samples from healthy and diseased individuals was performed as follows: i) discard protein groups present in less than 50% of the plasma samples within each study, ii) minimum value imputation and iii) statistical analysis using a two-sided Welch’s t-test (**Fig 4**). P-values were adjusted using the Benjamini-Hochberg procedure and proteins with FDR < 0.1 were considered significant. Pathway analysis was performed using Ingenuity Pathway Analysis (v81348237, Qiagen) and pathways with P-value < 0.05 were considered significant (**Fig. 4D**).

## ACKNOWEDGEMENTS

We would like to thank Jose Castro-Perez, Elliott Jones, Ihor Batruch, Patrick Pribil and Zoe Zhang from SCIEX for providing technical expertise and valuable comments on the manuscript and Vadim Demichev (Institute of Biochemistry - Charité) for guidance with DIA-NN. We are grateful to Kristina Kovacina, Kate Liu and Hui Yin Tan from AstraZeneca for sharing BAL and commercial plasma and sputum samples.

## AUTHOR CONTRIBUTIONS

KC, WS, YL, YH, AIR conceptualization; WS, YL, KC methodology; YL, WS investigation; YL, WS, KC validation; WS, KC data curation; KC, WS, YL formal analysis; KC, WS, YL visualization; KC software; YH, JC, ST resources; KC, AIR supervision; KC, WS, AIR project administration; KC, WS, AIR writing - original draft; KC, WS, AIR, YL, YH, JC, ST writing - review & editing.

## SUPPLEMENTAL DATA

This article contains supplemental data.

## SUPPLEMENTARY FIGURES

**Figure S1.**
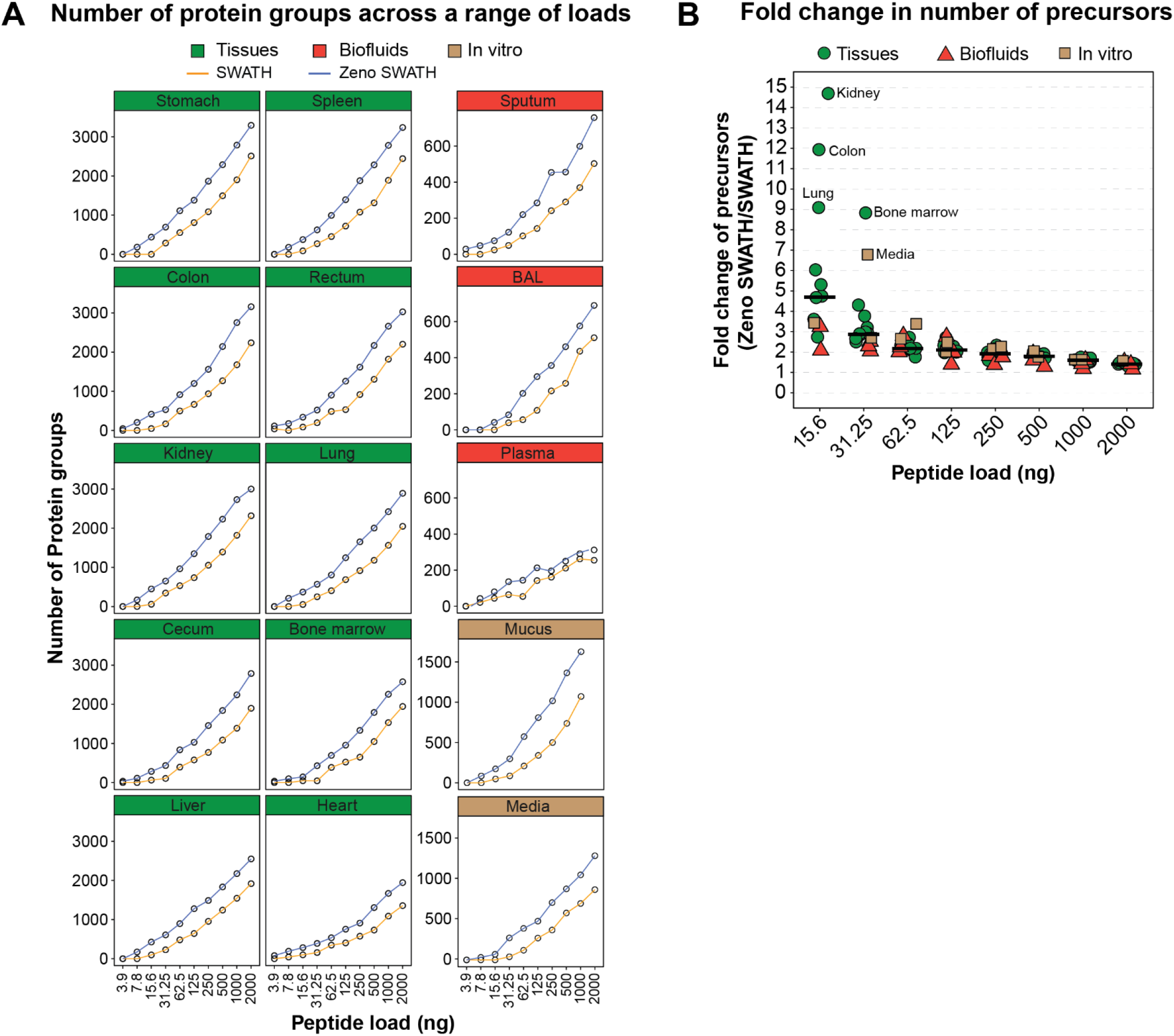
Sensitivity and protein coverage with Zeno SWATH DIA methods. (**A**) Number of protein groups identified across technical triplicates in various biological matrices at different peptide loads in SWATH and Zeno SWATH DIA modes. Peptide loads range from 3.9 ng to 2000 ng for all sample types except mucus that ranges from 3.9 ng to 1000 ng. (**B**) Fold change in precursors identified in Zeno SWATH DIA mode at different peptide loads. The horizontal bars represent the median.

**Figure S2.**
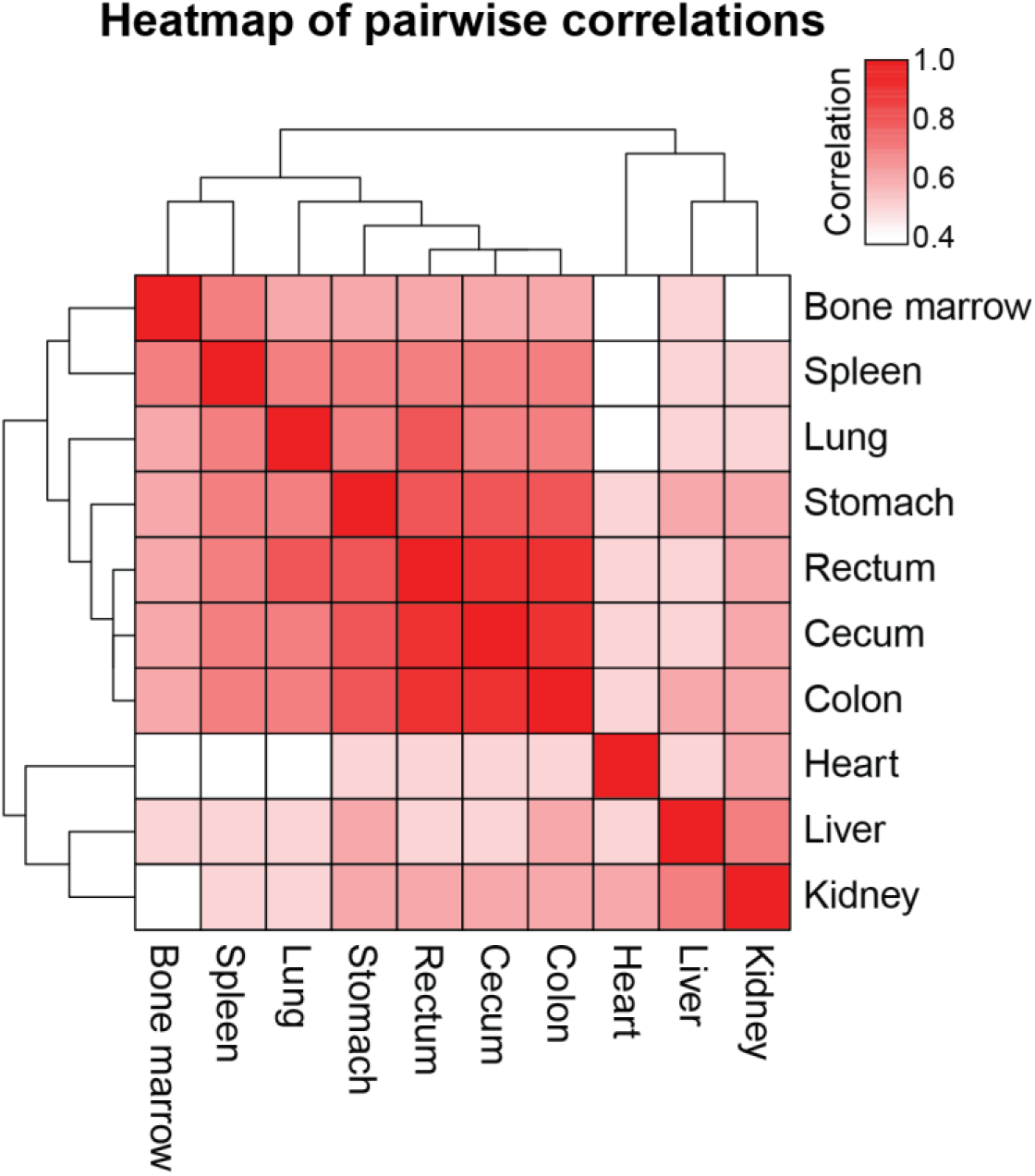
Similarity in protein composition across cynomolgus monkey tissues. Heatmap showing pairwise Pearson correlations between all 10 tissues. The median expression level across triplicates was used for each protein.

**Figure S3.**
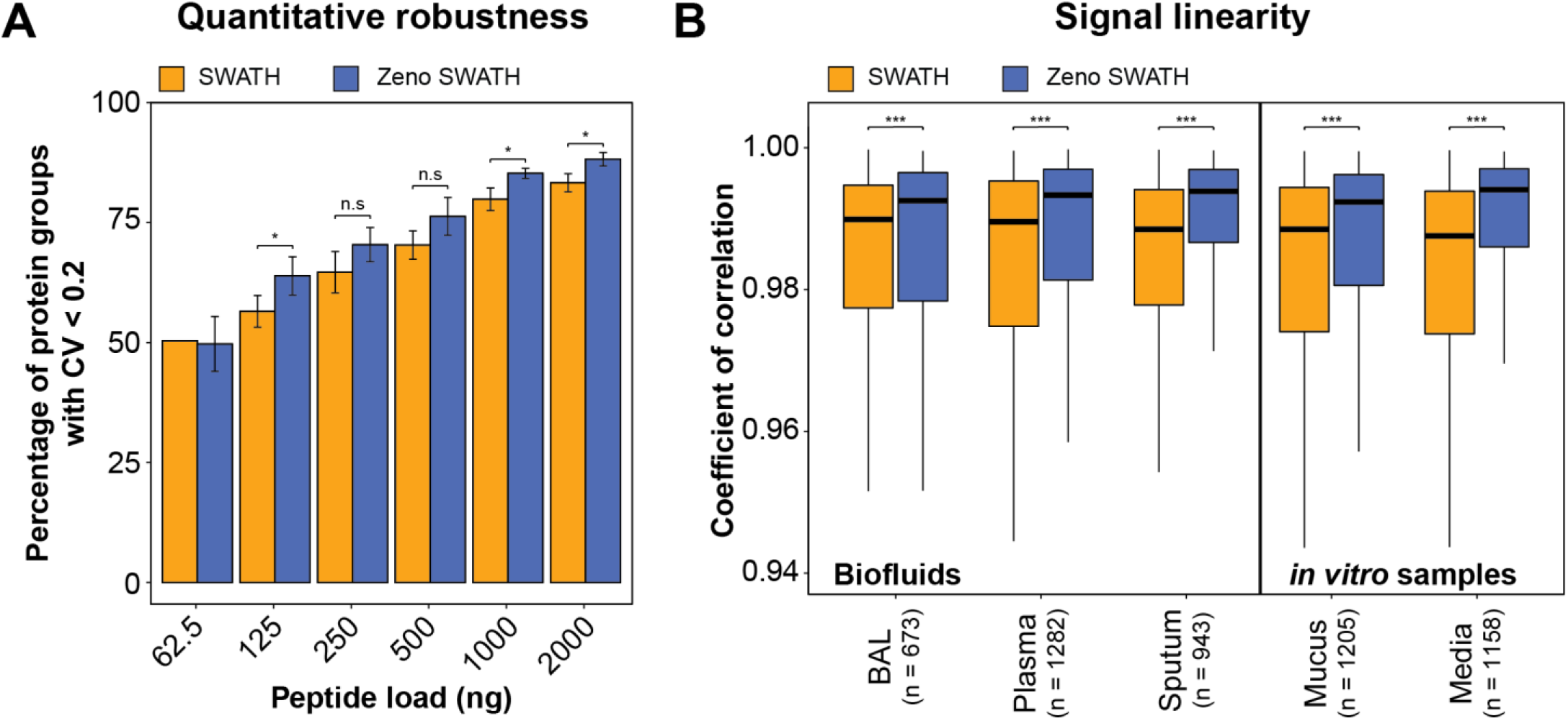
Evaluations of quantitative robustness and signal linearity in biofluid and *in vitro* samples. (**A**) Proportion of proteins with a coefficient of variation (CV) below 0.2 calculated across technical triplicates at different loads in SWATH and Zeno SWATH DIA modes. Only proteins detected in 100% of the triplicates were considered and only conditions with more than 50 proteins identified were plotted. Bars represent the mean across biofluid and *in vitro* samples and the error bars represent the standard deviation. A two-sided Welch’s t-test was used for differential analysis. n.s: not significant, *: P-value < 0.05, **: P-value < 0.01, *** P-value < 0.001. (**B**) Boxplot showing the distribution of Pearson coefficients of correlation of precursor abundances against dilution factor for biofluids and *in vitro* samples. Only precursors detected in at least 3 levels in both SWATH and Zeno SWATH DIA modes were considered in the analysis. The number of precursors is indicated in parenthesis. The box illustrates the first and third quartile, the whiskers are 1.5 times the interquartile range and the horizontal bar depicts the median of the distribution. A two-sided Mann–Whitney U test was used for differential analysis. *** P-value < 0.001.

## SUPPLEMENTARY TABLES

Table S1. Window number and width for SWATH and Zeno SWATH DIA experiments.

Table S2. Number of protein groups and precursors across LC-MS conditions for parameters optimization.

Table S3. Gain in pathway coverage across peptide loads and matrices with Zeno trap activated.

Table S4. Demographics information for healthy and diseased individuals.

Table S5. Differential proteins in plasma from patients diagnosed with NASH and T2D. Table S6. Differential pathways in patients diagnosed with NASH and T2D.

